# Modelling variation in bushmeat harvesting among seven African ecosystems using the Madingley Model: yield, survival and ecosystem impacts

**DOI:** 10.1101/695924

**Authors:** Tatsiana Barychka, Georgina M. Mace, Drew W. Purves

## Abstract

In principle, both the maximum sustainable yields of bushmeat, and the ecosystem impacts of extracting those yields, are likely to vary among ecosystems due to differences in the structure and function of ecosystems, but the data necessary to estimate this variation is lacking. Here, we compare seven different ecosystems on a North-South latitudinal gradient in Central Africa in terms of their trophic structure and capacity to support yields from bushmeat harvesting, using the Madingley General Ecosystem Model. The only factor that varies across simulations of these ecosystems is the climate which drives differences in vegetation structure and function, leading in turn to differences in the structure of the ecological community that emerge from the model. In a series of experiments (*n*=30), we simulate constant proportional harvesting of small and medium-sized warm-bloodied heterotrophs (1-23kg) over 30 years, recording expected bushmeat yields, and impacts on ecosystem structure, including trophic structure. Predictions for animal densities and trophic structures in the pristine (no harvesting) case varied among the ecosystems, with implications for bushmeat harvesting. For example, wooded savannah ecosystems stood out as having the greatest pristine densities in the target groups (11000-12000 animals per kilometre squared), greatest yields (100% higher than the tropical forest and 1000% higher than the desert ecosystem), and were the most resilient to harvesting. By contrast, small and medium-sized endothermic heterotrophs contributed only a small proportion of heterotrophs in the desert ecosystem, and thus the potential for bushmeat harvesting here was low. In all ecosystems, harvesting at the rate that maximised yield (55-65% population per year, except for the southern desert ecosystem) had strong impacts, causing drastic reductions in target functional groups, coupled with increases in smaller- and larger-bodied animals. Forest and desert ecosystems were particularly sensitive. Overall, the results suggest that, even for similar functional groups, bushmeat harvesting policies will need to vary substantially among ecosystems – and show that general ecosystem models could be a useful tool in helping to guide these policies.

## 1 Introduction

It has long been recognised that ecosystem structure and function, such as plant and animal biomasses, productivity, and turnover, are influenced by environmental conditions, including climate, soil quality and availability of water (Walter, 1964; Levin, 1998; Hunter and Price, 1992; Parrott and Meyer, 2012) - and Africa is no exception. Vegetation types have been linked to mean annual precipitation for a variety of ecosystems (Butt *et al.*, 2008; Del Grosso *et al.*, 2008; Hirota *et al.*, 2011), with almost linear relationships between primary production and rainfall reported by Whittaker (1970) and Walter (1964) in a range of African vegetation types. Clear empirical relationships between large herbivore biomass and mean annual rainfall have been described by Coe *et al*. (1976) in the east-African plains and savannahs, by Barnes and Lahm (1997) in central African forests, and by Bell (1982) in the woodland and savannahs of Africa. Similarly, in the tropical forests of Amazon and Guyana, Peres (2000) reported a positive relationship between primate biomass and soil fertility, where soil fertility was strongly correlated with annual rainfall.

Environmental correlations also exist at the species- and functional group levels, but these are not yet well documented for most species in Africa (though see Coe *et al*., 1976; McNaughton, 1976). Understanding of variation in ecosystem processes, and variation in interactions among species, is even less well developed, albeit improving (Hunter and Price, 1992; Montoya, Pimm and Solé, 2006). For example, food web models have been used to examine the role of various links within communities in maintaining their stability in the face of species removal (Sol and Montoya, 2001; Thompson *et al.*, 2012; Borrett *et al*., 2014). Nonetheless, the relationships between ecosystem structures and their responses to broad disturbances are still not well-understood (Montoya, Pimm and Solé, 2006). In addition, even the more complex food web models often ignore the environmental variability (Hunter and Price, 1992).

The variation in ecosystem structure and function across Africa implies that optimal harvesting policies and yields, as well as ecosystem impacts of harvesting are all likely to vary among ecosystems (Christensen and Walters, 2004; Fulton *et al.*, 2011; Mokany *et al.*, 2016). This in turn implies that the consequences of the dearth of data for guiding bushmeat harvesting are even more severe. In effect, a lot of data would be required to reliably estimate a good ‘one size fits all policy’ (even if such policy existed) for all of Africa. A lot *more* data would be required to find a whole set of such policies, tailored to the many different ecosystems where bushmeat hunting occurs. In addition, even if spatially and temporally reliable data on harvested species and ecosystems became available, predictions based on the empirical models (e.g. about sustainable harvest rates) would be specific to conditions and ecosystem responses described by the data (Boote, Jones and Pickering, 1996; Korzukhin, Ter-Mikaelian and Wagner, 1996), and would not account for the likely changes in biophysical conditions of the exploited systems, for example, due to climate change (Yates, Kittel and Cannon, 2000; Krinner *et al.*, 2005).

Species life history is certainly a key determinant for improving decisions about hunting efforts required for sustainable yields of bushmeat. However, in the absence of species-specific information, the use of a fully mechanistic ecosystem model: the Madingley Model (Harfoot *et al.*, 2014), could provide guidance, based solely on functional groups that are emergent from the ecosystem model (Purves *et al.*, 2013). The Madingley Model has been shown to give reliable predictions of trophic structure across a variety of terrestrial ecosystems (Harfoot *et al.*, 2014). The environmental inputs to the Madingley Model are spatially explicit, and include empirical data on air temperature, precipitation levels, number of frost days, seasonality of primary productivity and soil water availability (Purves *et al.*, 2013; Harfoot *et al.*, 2014). These inputs drive net primary production in the model. Plant and animal biomasses arise in the modelled ecosystems according to the locally specific climate and become components of ecosystem structure and function. Thus the distinctive feature of the model is that no species- or location-specific population parameters are input; they all emerge from the model structure and functions (Harfoot *et al.*, 2014). The Model has been used to explore independent and synergistic effects of habitat loss and fragmentation on ecosystem structure (Bartlett *et al.*, 2016), to predict non-linear regime shifts within ecological communities subjected to human removal of vegetation (Newbold *et al.*, 2018), to examine the importance of arbuscular mycorrhiza symbioses for the trophic structure of the Serengeti ecosystem (Stevens *et al.*, 2018) and to analyse properties of food webs (Flores *et al.*, 2019). Here, we use the model to simulate and compare dynamics of ecological communities that emerge in different ecosystems. Specifically, we use the Madingley Model to explore how maximum sustainable yields, optimum harvesting policy, and ecosystem impacts, might vary among different ecosystems – a question that is currently almost impossible to address using anything other than a general ecosystem model. We are modelling species populations for which no population parameters are available, but whose dynamics are determined entirely by the ecosystem model.

We use the Madingley Model to simulate the effect of constant proportional harvesting of small and medium-sized heterotrophs in seven ecosystems on a North-South latitudinal gradient through Central Africa. The objective is to compare how different harvesting levels drive ecosystem changes, ecological community structure and productivity. By only varying the model’s environmental inputs while keeping all the other model inputs (such as the starting number of cohorts and stocks, and harvest rates) constant (Figure 1), any differences between ecosystems that result from harvesting are attributable to differences in ecosystem structure and functioning, as predicted by the model.

**Figure 1.**
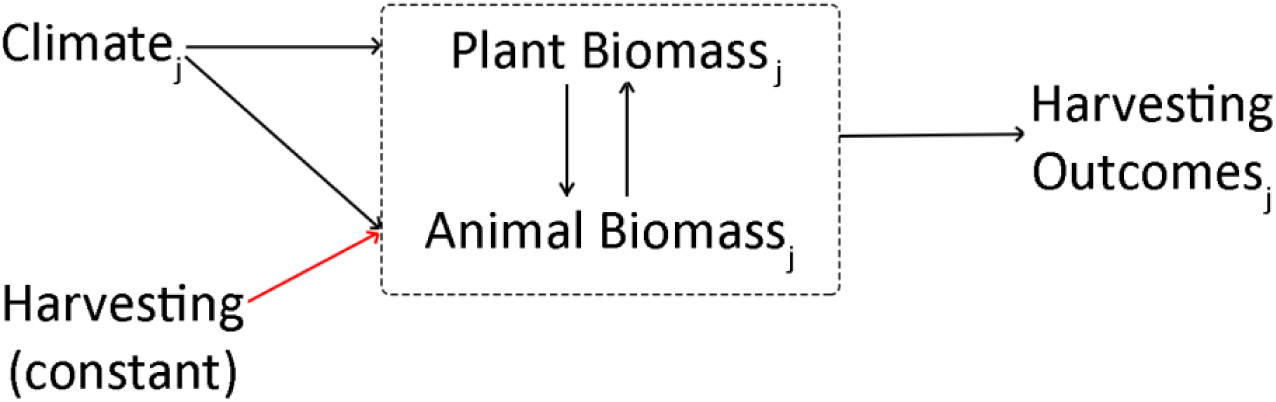
Summary of the experiment: climate for *j*-grid cells (n=7) is input into the Madingley Model; ecosystem structure emerges as a result of climate and of multi-trophic interactions; the same level of proportional harvesting is applied to all *j*-ecosystems; harvesting outcomes are output for each ecosystem *j* from 1 to 7.

Our expectation is to see marked differences in expected bushmeat yields and in sensitivity to harvesting between ecosystems. If successful, these experiments could contribute to the debate about the importance of environmental conditions in predicting ecosystems dynamics, and about the potential of large-scale models such as the Madingley Model, for supporting land-use and conservation policies.

## 2 Methods

### 2.1 The Model

The Madingley Model: a) receives environmental data based on user-defined latitude and longitude: location-specific empirical data on air temperature, precipitation levels, number of frost days, seasonality of primary productivity and soil water availability; b) predicts ecosystem dynamics from environmental inputs, and animal and plant dynamics described in the model using a set of core biological and ecological processes (plant growth and mortality, and eating, metabolism, growth, reproduction, dispersal, and mortality for animals); and c) outputs estimates of biological characteristics of the emergent ecosystem (Harfoot *et al.*, 2014).

The Madingley Model represents the state of the animal part of the ecosystem in terms of the densities of individual animals with different functional traits. The densities change through time as individuals interact, in turn resulting in births, deaths, growth rates, and dispersal, with the interactions (e.g. predation) defined entirely in terms of the functional traits. Although the model is defined entirely in terms of interactions among individuals, the simulation uses a computational approximation (based around so-called cohorts) to allow for all interactions among all individuals to be simulated. The animal part of the ecosystem is ultimately fed by the vegetation, which is simulated using a simple stock and flow model, driven by climate, but affected by herbivory. For detailed description of the Model see Harfoot *et al.* (2014).

### 2.2 Locations

Seven locations on a North-South latitudinal gradient in Africa were selected (Figure 2) to represent seven ecosystems in three broad vegetation types (Otte and Chilonda, 2002): savannah (grass and shrub, and wooded savannah in the North and South), forest (tropical forest, and woodland and shrub), and desert (North and South). Each ecosystem was modelled by a one-degree geographic grid cell (approximately 12307km^2^), centred on the coordinates provided in Appendix 1. No inter-cell migration was included in the simulations.

**Figure 2.**
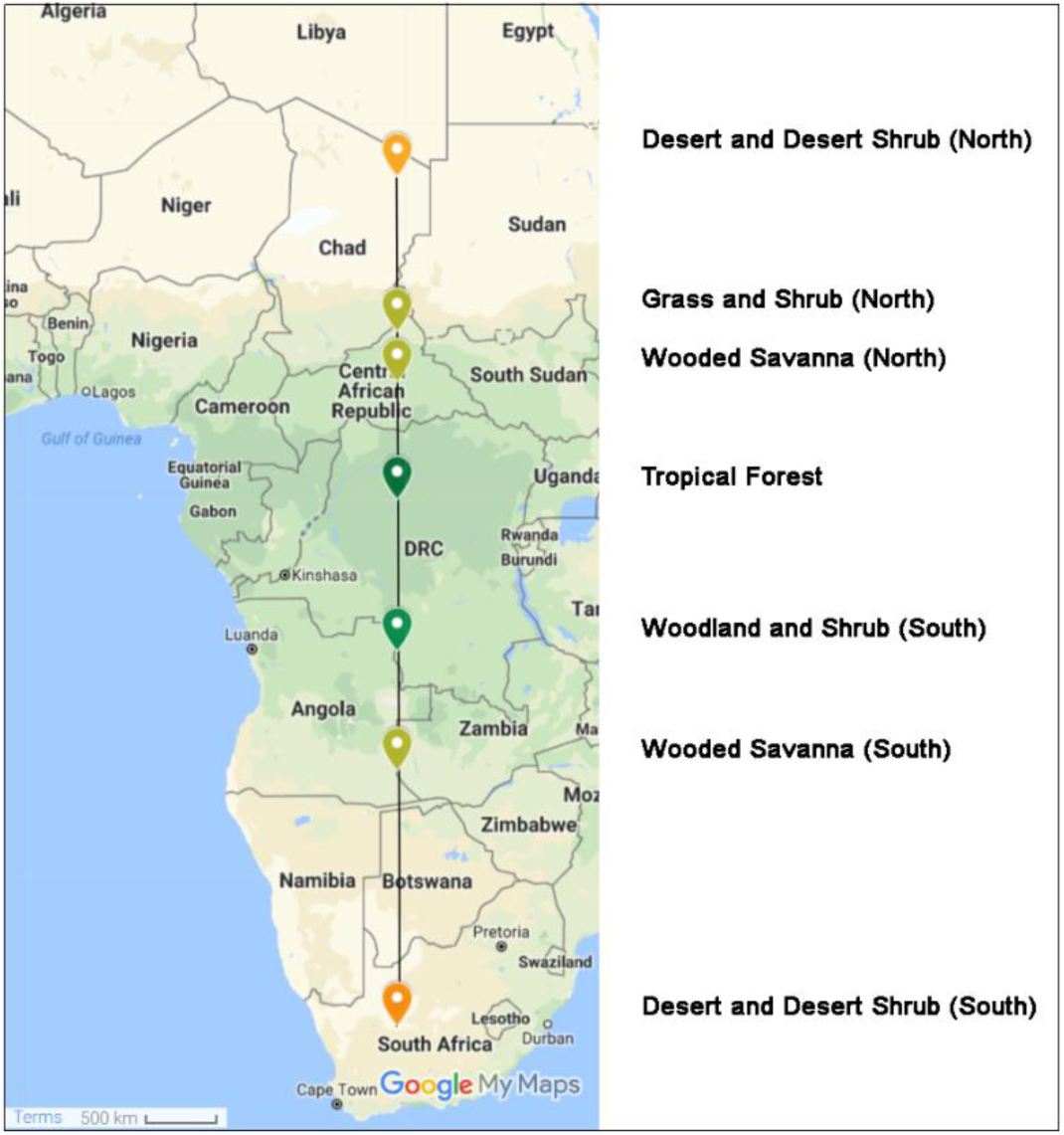
The locations of sites used for harvesting simulations, representing seven ecosystems in three broad vegetation types: desert (orange), savannah (light green) and forest (dark green).

### 2.3 Harvesting simulations

In all sites the same harvesting strategy was applied targeting small and medium-sized endothermic carnivores, omnivores and herbivores with adult body masses of between 1kg and 23kg, and over 100 grams as juveniles, based on reported bushmeat species sizes in Afrotropical forests (Fa, Ryan and Bell, 2005).

In each ecosystem, constant proportional harvesting was applied to target animals for 30 years. All harvesting took place once a year in a single month (set at month 6) to approximate discrete harvesting used in most conventional bushmeat harvesting models. The harvest rate (*φ*) was set to 0 ≤ *φ* ≤ 0.90 in increments of 0.05 from 0.0 ≤ *φ* ≤ 0.70, and in increments of 0.10 thereafter. We used smaller rate increments for *φ* between 0 and 0.70 to get more detailed estimates of harvesting outcomes (yield, survival and ecosystem impacts) at moderate-to-high harvest rates.

Each harvesting scenario was simulated 30 times (*n=30*) for each ecosystem; i.e. 17 harvest rates per ecosystem were replicated 30 times resulting in 510 model runs for each location and 3570 model runs in total.

For each ecosystem simulation, a 1000-year burn-in (*n=30*) was run. Ecological communities were allowed to emerge in the model and to reach equilibrium states in terms of number of different cohort types (animal body mass, herbivore/omnivore/carnivore, and ectotherm/endotherm). These states were then used as the initial state for harvesting simulations in the given location.

The recorded ecosystem states without harvesting (simulations where *φ*=0) are the reference for the ‘pristine state’ of the ecosystem.

### 2.4 Outputs and processing

#### 2.4.1 Trophic Pyramids

For each location, all heterotrophs were identified as ectotherms or endotherms belonging to one of the three functional groups (carnivores, omnivores or herbivores). Individuals in each functional group were also allocated into a body mass bin (*b*), ranging from the smallest (0.1-0.3kg) to the largest body size (316.2-1000kg). The smallest body mass bin (*b* = −2) ranged from: 10^−2^ to 10^−1^ gram; and the largest bin (*b* = 6) ranged from: 10^6^ to 10^7^ gram. Because some of the bins were deemed too wide to be able to capture changes in cohort abundances due to harvesting, bins were further sub-divided into smaller sub-bins, where adult body masses were incremented in steps of 0.5 for 2 ≤ *b* ≤ 6 (e.g. 10^3^-10^3.5^ gram body mass bin contains animals 1-3.2kg in size).

To examine the trophic structure of pristine ecosystems, the total biomasses in each functional group and body mass bin in the final year of each simulation were summed, and then averaged across the 30 replicates.

#### 2.4.2 Harvesting Outcomes

For target individuals only, total yields from harvesting and population densities were recorded during harvesting in years 0-30. For all individuals (target and non-target), we recorded ecosystem-level information such as: functional group identifiers, abundances, and adult and individual body masses, in years 0, 10, 20 and 30.

Using total population densities, we calculated the probability of persistence of the target animals, assuming that a 90% and a 99% reduction in total population density (compared to the pristine density in month 0, after the burn-in) at any point during the simulation run constituted a high risk and a very high risk of extinction, respectively (Mace and Lande, 1991). Each simulation run was assigned a one or a zero depending on whether total population densities did (0) or did not (1) decline by 90%/99% at any point during the simulation run. The outcomes were averaged across simulations to give an estimate of animal persistence for each harvest rate, by location.

We define harvesting levels which could result in a high probability of extinction (declines of 90%) in at least 10% of the cases (i.e. 10% of simulations) as the high risk harvesting, and harvesting which could result in very high probability of extinction (declines of 99%) in at least 10% of the cases (i.e. 10% of simulations) as the very high risk harvesting.

We define three harvesting strategies:

*Maximum harvesting* strategy/maximum harvest rate – harvesting that maximises yield over 30 years.

*Constrained high risk* strategy – harvesting that maximises yield over 30 years, subject to the constraint of high risk of extinction (i.e. harvest rates are constrained to ensure at least 10% of population survive on average in at least 90% of the cases).

*Constrained very high risk* strategy – harvesting that maximises yield over 30 years, subject to the constraint of very high risk of extinction (i.e. harvest rates are constrained to ensure at least 1% of population survive on average in at least 90% of the cases).

To examine the potential effects of harvesting, we measured changes in abundances of animals in different body mass bins at different levels of harvesting pressure, focusing on the group directly impacted by harvesting: the endothermic heterotrophs.

All data processing, statistical analysis and visualisation were done in R version 3.5.1 (R Core Team 2018), with minor editing (image stitching and adding text to images) in Adobe Photoshop CC.

#### 2.4.3 Per Capita Yield Conversion

In order to compare bushmeat yield to that of farmed cattle in the same ecosystems, we collected estimates of human population density and beef offtakes by agro-ecological zones from Otte and Chilonda (2002); human population density estimates were used to convert bushmeat yields per kilometre squared to bushmeat yields per kilometre squared per capita.

## 3 Results

### 3.1 Trophic structure of modelled ecosystems

The total heterotroph biomasses by functional group, and by functional group and body size in seven pristine ecosystems are presented in Figure 3 and Figure 4, respectively. The highest heterotroph biomasses were in the savannah (2.4-3.1 million tonnes) and forest ecosystems (2.3-2.6 million tonnes), followed by desert and desert shrub in the South (2.1 million tonnes) (Figure 3). Only around 1% (0.1 million tonne) of total heterotroph biomass was present in the northern desert (Figure 3).

**Figure 3.**
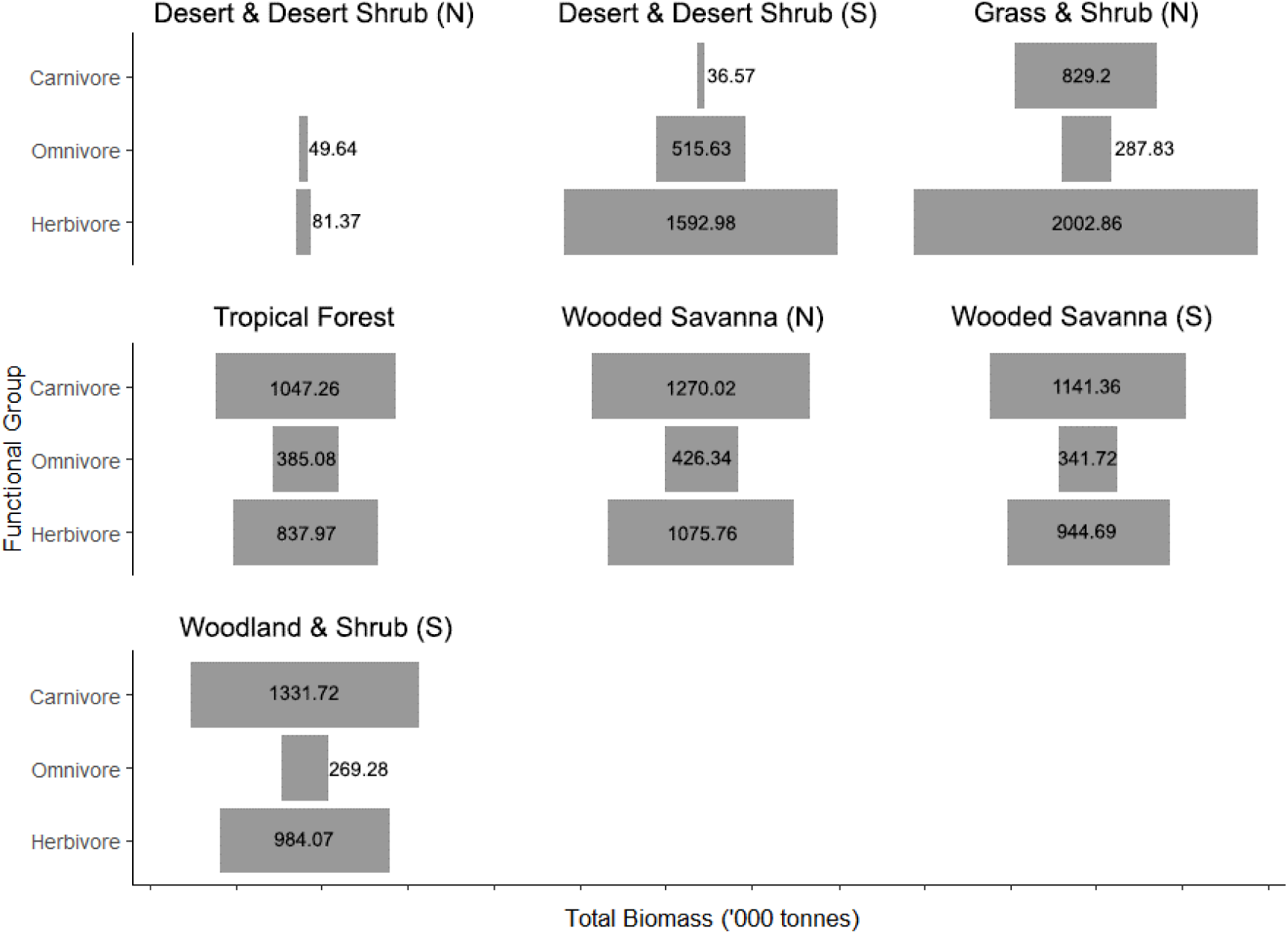
Trophic biomass pyramids in seven pristine ecosystems. Numbers inside or next to the bars represent total endotherm and ectotherm biomass (‘000 tonnes).

**Figure 4.**
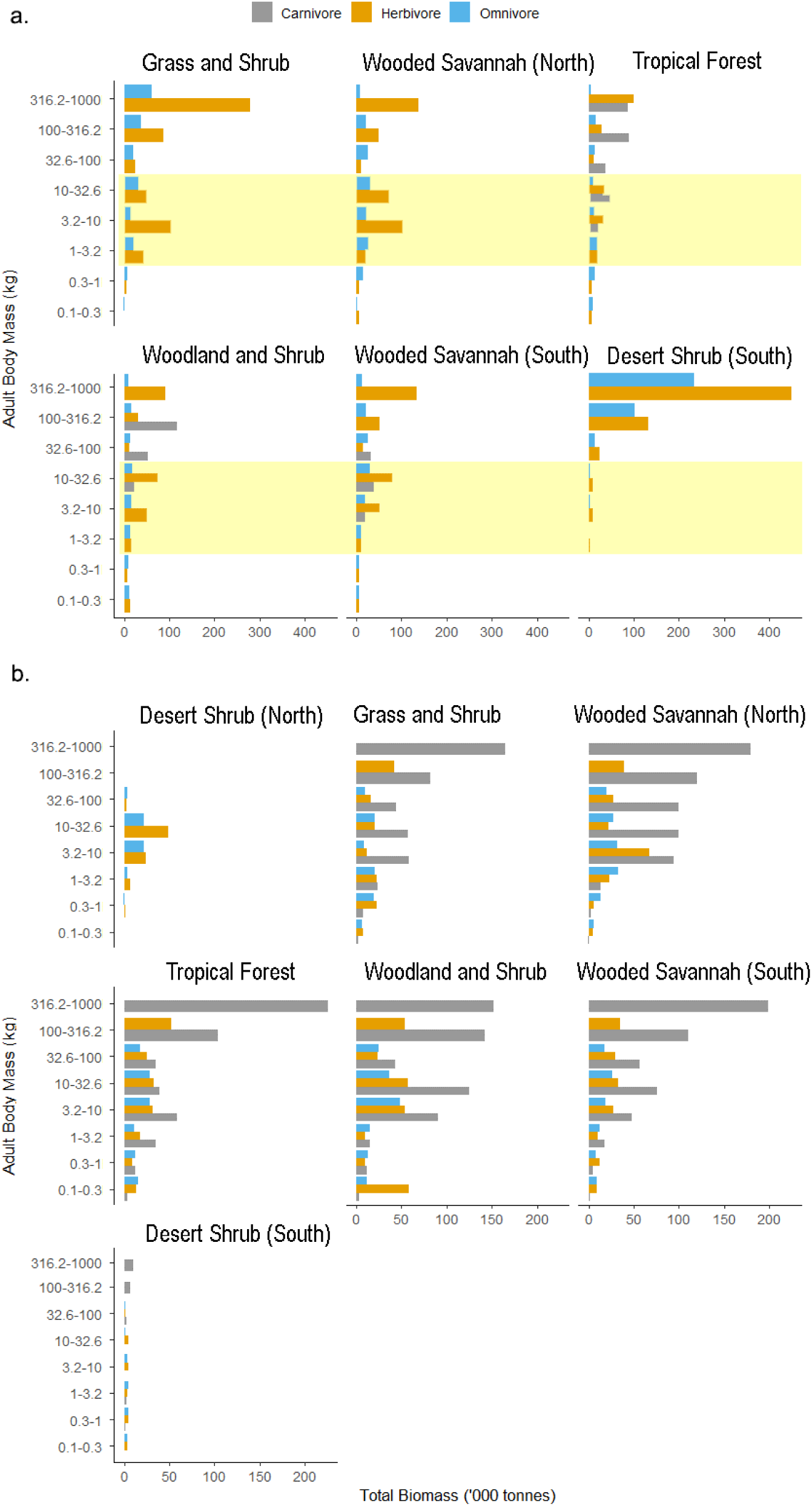
Total biomass (‘000 tonnes) of endothermic (a.) and ectothermic (b.) heterotrophs (carnivores, omnivores and herbivores) in different body mass bins in pristine ecosystems, by ecosystem. Targeted populations are indicated by yellow band in a. For clarity, very large (>1000kg) and very small (<0.1 kg) organisms had been removed.

The carnivores (Figure 3 and grey bars in Figure 4) were the dominant functional group in the forests (between 46%-52% of average total biomass in these ecosystems) and wooded savannah ecosystems (46%-47% of average total biomass in wooded savannah), but not in the grasslands (27%) or desert ecosystems (0%-2%). The herbivore (Figure 3 and orange bars in Figure 4) contribution to total biomasses was the highest in the desert ecosystems (62%-74% of average total biomasses) and the grasslands (64%), and the lowest in the forests (37%-38%). The omnivores (Figure 3 and blue bars in Figure 4) had the lowest total biomasses of all functional groups in all productive ecosystems (9%-17% of average total biomasses). In the deserts, the omnivores had the second-highest biomass densities after the herbivores (24%-38% of the total heterotroph biomasses).

In terms of body-size composition (Figure 4), all ecosystems had relatively high proportion of total ecosystem biomasses represented by top predators, i.e. large-bodied (>316.2kg) carnivores (around 40% of all carnivores in the forest and desert ecosystems, and 26%-30% in the wooded savannah ecosystems). The lowest biomass proportion of large carnivores was in the southern desert (approximately 1% of the total biomass), which coincided with the highest biomass proportion of large-bodied (100-316.2kg) herbivores. Interestingly, the model predicted the carnivores to be predominantly ectothermic (e.g. 99%-100% of the total carnivore biomasses in the savannah and the southern desert, on average), even in the larger body mass bins. The highest share of endothermic carnivores was in the tropical forest (around 11% of estimated total carnivore biomass in that ecosystem). The highest total biomasses of large-bodied (>100kg) endothermic herbivores were in the southern desert and in the grasslands ecosystem (Figure 4a): 13% and 7% of the total biomasses in these ecosystems, respectively, vs around 3% of the total biomass in the tropical forest ecosystem.

Targeted small and medium-sized endothermic herbivores (Figure 4a, highlighted by the yellow band) had the highest biomasses in the northern savannah ecosystems (21-103 thousand tonnes), followed by the southern wooded savannah and woodland and shrub (30% lower than the northern savannah, on average; 11-79 thousand tonnes), the tropical forest (on average, 55% lower than the northern savannah; 20-35 thousand tonnes), and the southern desert ecosystem (90% lower than the northern savannah; 3-10 thousand tonnes). Similarly, for small and medium-sized endothermic omnivores (Figure 4a, highlighted by the yellow band), the highest total biomasses (15-33 thousand tonnes) were returned in the northern savannah ecosystems, followed by the southern savannah (10-29 thousand tonnes) and forest ecosystems (12-19 million tonne; 30%-40% lower than the northern savannah), and the southern desert (circa 2 thousand tonnes; 90% lower than the northern savannah). Small and medium-sized (1-32.6kg) warm-bloodied carnivores were only present in the forests and in the wooded savannah in the South (22-50 thousand tonnes, and 20-39 thousand tonnes, respectively).

Endothermic heterotrophs were entirely absent from the northern desert ecosystem (Figure 4) (hence no bushmeat harvesting was modelled in the northern desert, see Results). The northern desert was dominated by small and medium-sized (3.2-32.6kg) ectothermic omnivores and herbivores.

### 3.2 Harvesting Outcomes

#### 3.2.1 Population persistence

Qualitatively, targeted populations’ responses to harvesting were similar between ecosystems (Figure 5). The proportion of persistent populations decreased with harvesting intensity, and, with the exception of the grassland and shrub, for each level of risk (high vs very high), there existed a threshold, beyond which persistence over 30 years declined rapidly. At harvesting rates below the threshold, persistence was high, and showed no relationship with harvest rate, or only a slight relationship. At harvest rates above the threshold, persistence declined rapidly with increasing harvesting. The exception was in the grassland ecosystem, for the high risk case. Here, persistence with no harvesting over 30 years was significantly lower than in the other ecosystems (47%±18%; 95%CI, *n*=30, compared to 83-100% in all other ecosystems), and the relationship between persistence was closer to linear, such that persistence declined steadily with increasing harvesting over the full range of harvesting rates. For the remaining cases, despite the general qualitative agreement, the location of the thresholds (i.e. the harvesting rates that caused persistence probabilities to begin to rapidly decline) differed between locations and according to the level of risk. The thresholds were closer for the two wooded savannah ecosystems (circa 0.45 vs circa 0.60), and for the two forest ecosystems (circa 0.25 vs circa 0.20). For all ecosystems, there were marked differences between the risk cases (the high vs the very high risk case; the orange and the green line in Figure 5), with the highest discrepancy between trajectories in the forests, and the grassland and shrub.

**Figure 5.**
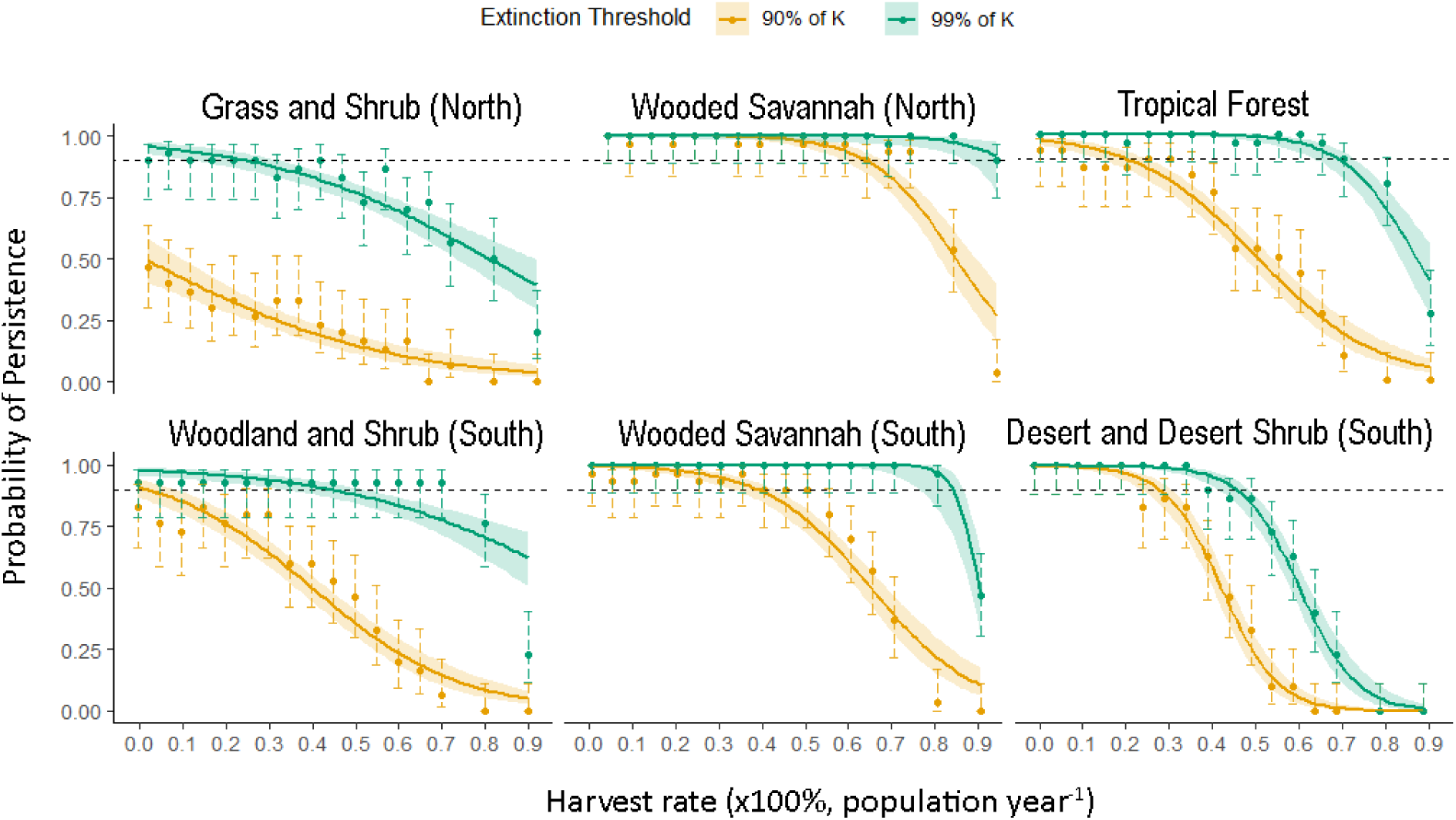
Probability of target species persistence (with 95% confidence interval in shaded orange/green, and 2 standard errors shown with error bars) with harvesting intensity over 30 years. On y-axis, values close to 1 indicate population density declining by 90% (in orange) or 99% (in green) during the 30-year harvesting period in only few replicate simulations; values below the horizontal dashed line indicate populations decline by 90% and 99%, respectively, in over 10% of the cases (high and very high risk of extinction, respectively) over 30 years.

For setting real-life harvesting policies, the thresholds could, in principle, be used to set the maximum allowable harvesting rate. Of all ecosystems, wooded savannah was the most resilient to harvesting according to this metric, with a potential to accommodate harvest rates of up to 40%-60% population year^−1^ under the constrained high risk strategy (at least 10% of initial population survived on average in at least 90% of the cases, corresponding to the portion of the orange trend lines above the horizontal dashed line in Figure 5). These ecosystems also supported the highest densities of small and medium-sized heterotrophs in the pristine state (approximately 11000-12000 animals km^−2^, Appendix 2). Here, the target population density declined by 4%-6% for each 5% increase in effort up to the annual harvest rate of 70% of population, with 26%-28% drop in density per 10% increase in effort thereafter (Appendix 3).

In the tropical forest, the thresholds were much lower than in the wooded savannah, allowing harvesting up to around 20-30% population year^−1^ under the constrained high risk strategy. Pristine population density of small and medium-sized heterotrophs was lower compared to the wooded savannah (around 9000 animals km^−2^; Appendix 2); and the average densities declined by 6% for each 5% increase in effort up to 70% population per year (Appendix 3). In the southern desert, the maximum allowable harvest rate was 20%-25% population year^−1^ under the constrained high risk strategy, with the estimated target population densities of approximately 2700 animals km^−2^ (Appendix 2), declining by 16% for each 5% increase in effort up to 70% population per year, and by 37%, thereafter (Appendix 3).

In the woodland and shrub and the grassland ecosystems, the background extinction rates were above 10% of population on average. It was therefore impossible to find any harvest rates that returned a population persistence above 90%. Therefore, harvesting was only feasible under the constrained very high risk strategy, i.e. accepting that up to 99% of population could be lost due to a combination of harvesting and natural mortality (the green trend line in Figure 5). The percentage declines in average density of target populations with harvesting were lower in the grasslands than in other ecosystems: by 4% for each 5% increase in effort up to the annual harvest rate of 70% population per year, and by 12%, thereafter (Appendix 3).

#### 3.2.2 Meat Yields

The yields returned by the maximum, vs the constrained high risk, harvesting of small and medium-sized heterotrophs varied substantially among the ecosystems (Table 1). The average yield varied widely across ecosystems. In the wooded savannah, the maximum yield was almost twice that of forest ecosystems, over 200% higher than in the grassland and shrub, and almost 1000% higher than yields in the desert ecosystem (Figure 6). Yields in the grasslands and the tropical forest were comparable; however, the probability of low yields was significantly higher in the grassland and shrub ecosystem (note strong right skew in Figure 6).

**Table 1.** Bushmeat yields under the maximum and the constrained high risk strategies (in kg km^−2^ year^−1^ and in kg capita^−1^ year^−1^, with 1 standard error, s.e.) by ecosystem, compared to the empirical beef offtakes (in kg capita^−1^ year^−1^).

| Ecosystem | Bushmeat |  |  |  | Beef<br>Offtake,<br>kg capita<sup>-1</sup><br>year<sup>-1</sup> |
| --- | --- | --- | --- | --- | --- |
|  | Yields, kg km<sup>-2</sup> year<sup>-1</sup><br>(mean±s.e.) |  | Yields, kg capita<sup>-1</sup> year<sup>-1</sup><br>(mean±s.e.) |  |  |
|  | Maximum | High risk | Maximum | High risk |  |
| Grass and<br>Shrub | 2221.73<br>(±83.43) | - | 67.53<br>(±2.54) | - | 11.1 |
| Wooded<br>Savannah<br>(North) | 6722.15<br>(±142.45) | 6722.15<br>(±142.45) | 652.64<br>(±13.83) | 652.64<br>(±13.83) | 7 |
| Tropical<br>Forest | 3407.44<br>(±68.00) | 2246.98<br>(±36.89) | 224.17<br>(±4.47) | 147.83<br>(±2.43) | 1.9 |
| Woodland<br>and Shrub | 3675.97<br>(±94.16) | - | 356.89<br>(±9.14) | - | 7 |
| Wooded<br>Savannah<br>(South) | 6319.67<br>(±126.45) | 5635.69<br>(±105.54) | 613.56<br>(±12.28) | 547.15<br>(±10.25) | 7 |
| Desert and<br>desert shrub<br>(South) | 616.97<br>(±11.49) | 497.88<br>(±6.62) | - | - | - |

**Figure 6.**
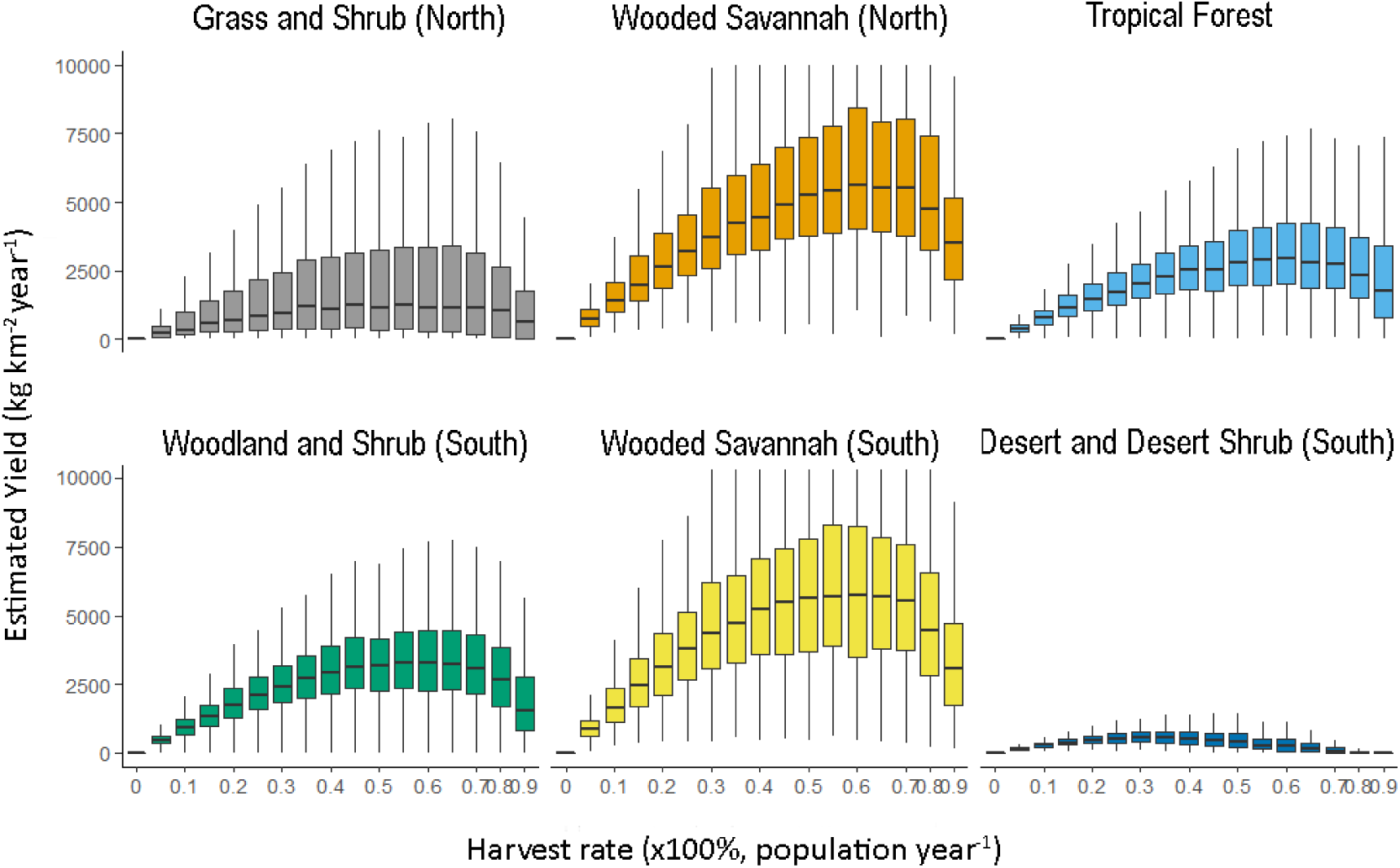
Average meat yields with harvesting intensity (not constrained by probability of persistence), by ecosystem.

The harvest rate that maximised yield (the maximum rate, Table 2) was 55%-65% population year^−1^ in all ecosystems, except for the desert and desert shrub (around 30-35%). In all ecosystems, harvesting at the maximum rate reduced target population densities by at least 90%, compared to their pre-harvest densities (i.e. high or very high risk of extinction). In wooded savannah ecosystems and southern desert, the maximum rates and the constrained high risk rates were similar within ecosystems (Table 2). In the grasslands and in both forest ecosystems, the maximum harvest rates were significantly higher than harvest rates under the constrained high risk (but not the very high risk strategy); for example, 60% vs 30% population year^−1^ in the tropical forest (Table 2). I.e. in the grasslands and forest ecosystems, maximising yield could result in extinction of 90-99% of animal population.

**Table 2.**
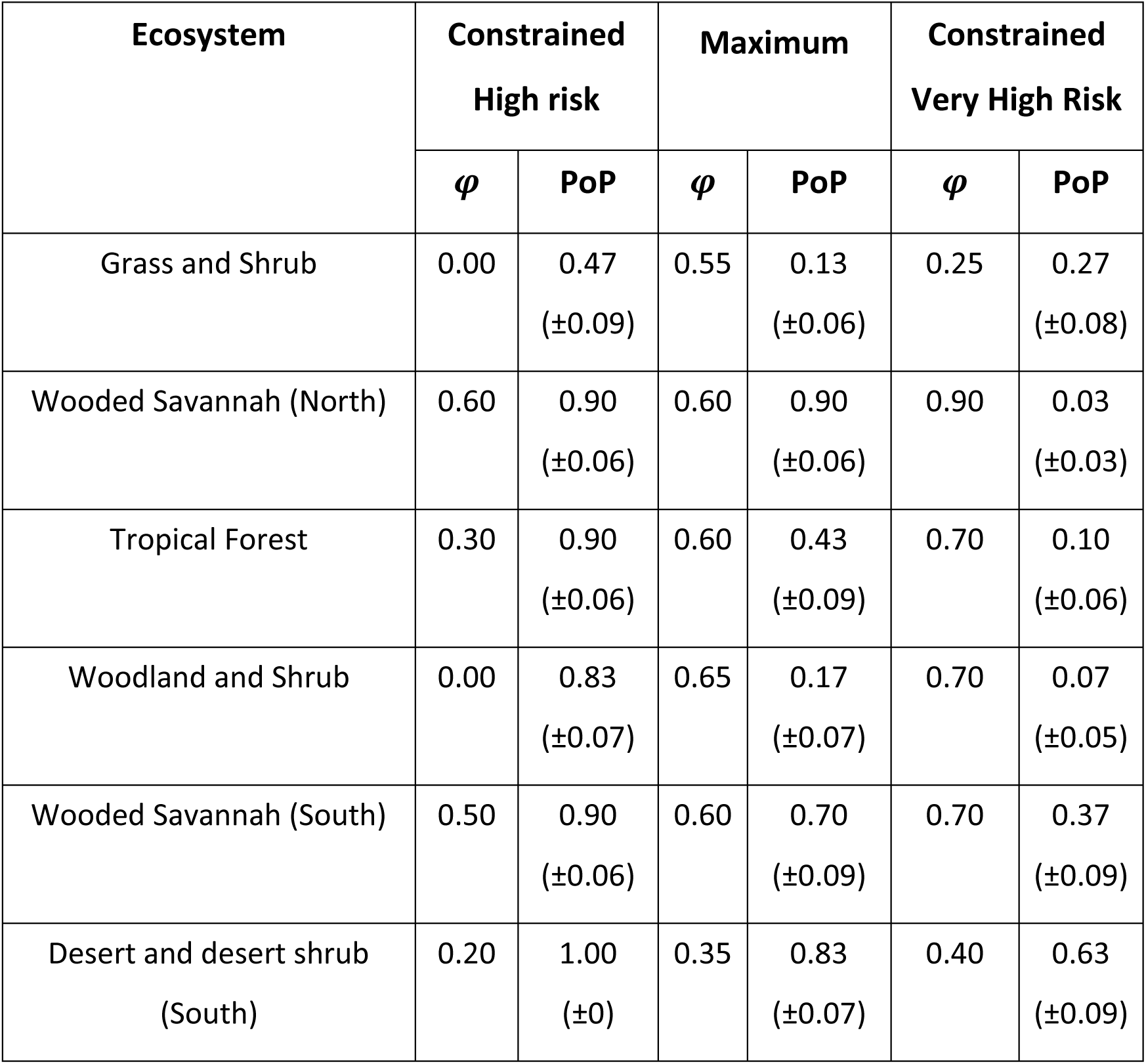
Harvest rate, *φ* and associated probability of persistence, PoP (calculated for the high risk harvesting, orange line in Figure 5-5; ±1 standard error, 95% CI, *n*=30) over 30 years, by harvesting strategy (constrained high and very high risk, and unconstrained maximum harvesting), by ecosystem.

In all ecosystems bar one (the grass and shrub ecosystem), the maximum harvest rates were below the harvest rates under the constrained very high risk strategy (Table 2). The corresponding yields were the opposite: the maximum yields were above the yields under the very high risk harvesting (Figure 6). This suggests that using the 1% survival threshold to set harvest rates (the very high risk strategy) was sub-optimal compared to the maximum harvesting in terms of species survival and in terms of meat yields.

Average bushmeat yields per capita per year under the maximum harvesting strategy were 6-117 times higher than the beef offtakes in sub-Saharan Africa, with the smallest difference (6 times) in the grasslands and the highest (117 times) in the tropical forest (Table 1); however, maximum harvesting was associated with high risk of extinction in all ecosystems except for wooded savannah in the North (Table 2). An estimate of human population density for the southern desert ecosystem was not available (possibly, very low); therefore, we couldn’t calculate per capita bushmeat yields.

#### 3.2.3 Impacts of harvesting

Across the ecosystems, and considering harvesting at three levels of intensity (20%, the maximum rate for each ecosystem, and 90%), there was evidence of a shared pattern of responses to harvesting, compared to the pristine baseline (Figure 7 and Figure 8). First, functional groups targeted for harvesting, i.e. mid-sized (1-23kg) herbivores, omnivores, and carnivores, tended to decline, as might be expected given that they were being removed. Second, within the target functional groups, omnivores tended to decline more than herbivores. Third, within the target functional groups, larger-bodied herbivores tended to decline more than smaller-bodied functional groups. Fourth, the declines in the targeted functional groups were coupled with increases in smaller-bodied non-targeted herbivores and omnivores, and less pronounced increases in larger-bodied non-targeted herbivores and omnivores. There were exceptions to this general pattern, and the individual changes were often not statistically significant. Nonetheless, comparing all responses together, the overall pattern was relatively clear (Figure 7 and Figure 8). However, there were marked differences in responses to harvesting between ecosystems, and between the functional groups within ecosystems.

**Figure 7.**
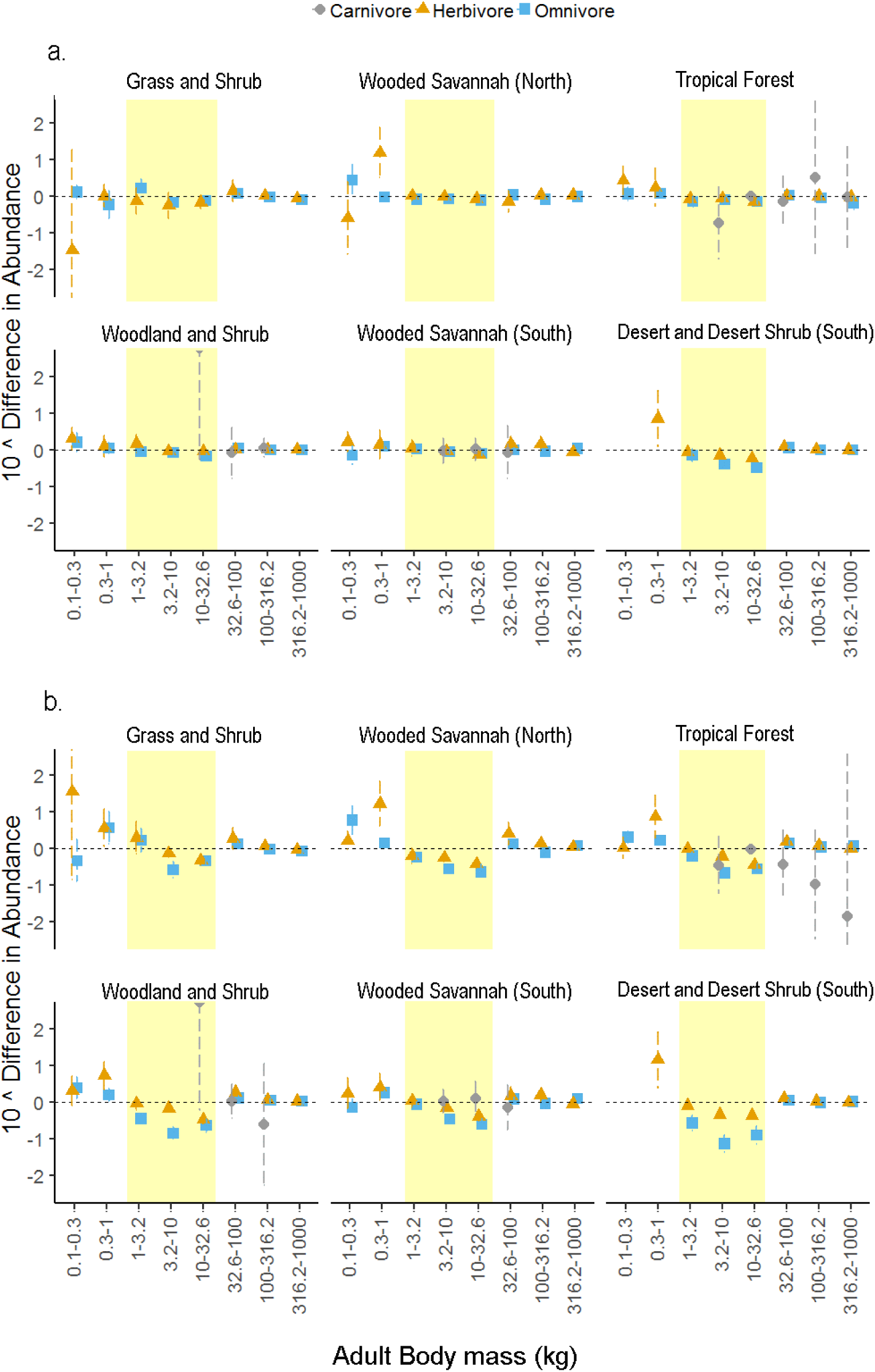
Changes in abundances of endothermic heterotrophs (with 95% confidence intervals) as a result of harvesting small-to-medium sized heterotrophs (highlighted in yellow) at the rate of 20% of population year^−1^ (in a.), and at the maximum rate of harvesting (in b.), by ecosystem and adult body mass. The horizontal dashed line indicates no significant impact of harvesting on abundances.

**Figure 8.**
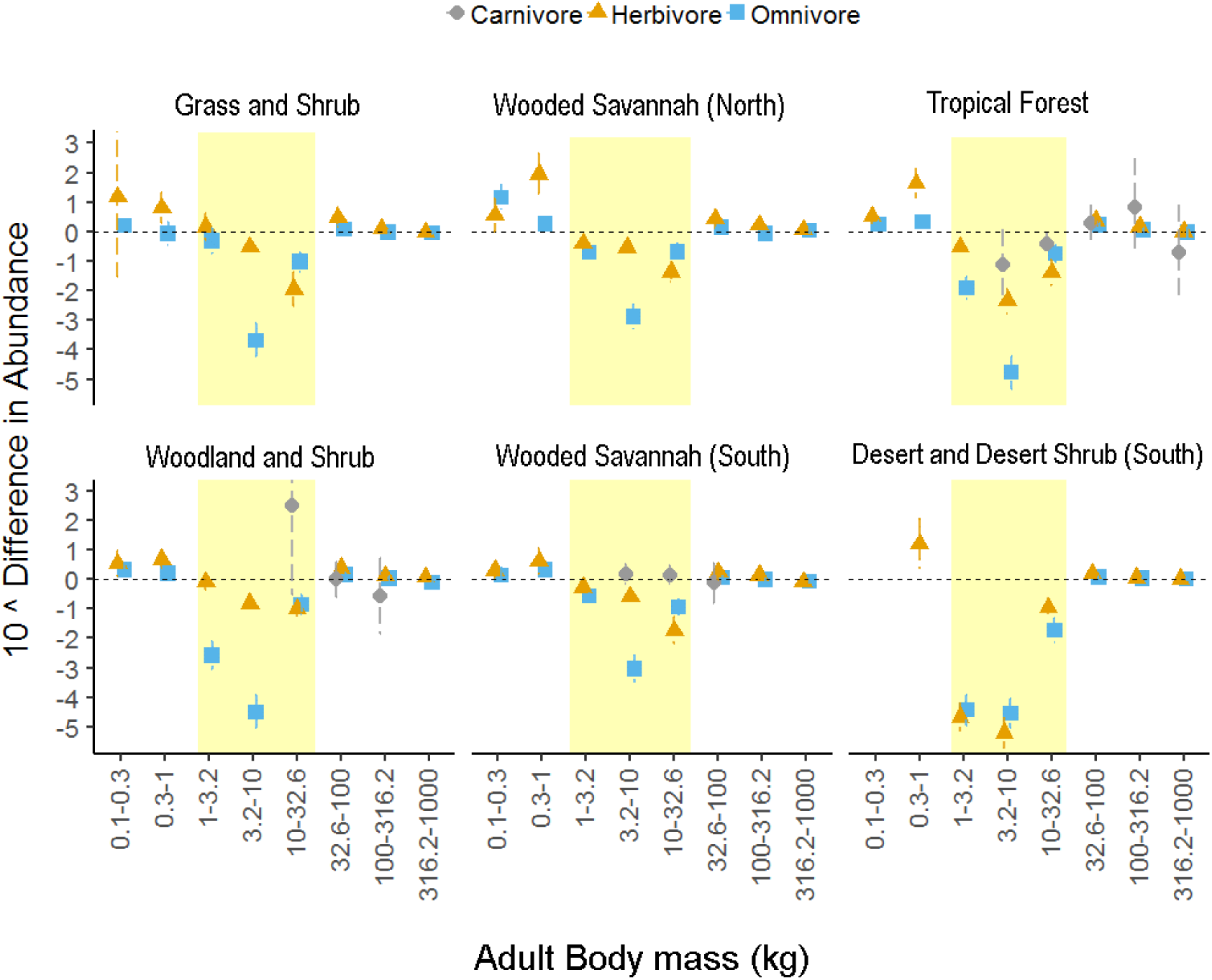
Changes in abundances (with 95% confidence intervals) of endothermic heterotrophs (with 95% confidence intervals) as a result of harvesting small-to-medium sized heterotrophs (highlighted in yellow) at the rate of 90% of population year^−1^, by ecosystem and adult body mass. The horizontal dashed line indicates no significant impact of harvesting on abundances.

Harvesting 20% of population per year had no statistically significant impact on target cohorts in any of the ecosystems (Figure 7a), except for the southern desert. Here, densities of omnivores and medium-sized (3.2-32.6kg) herbivores declined by 58%-66% and by 27%-39% on average, respectively. Non-target small-bodied (<1kg) herbivores and omnivores became more abundant in all ecosystems; however, at this level of harvesting, the effect of harvesting on small-bodied heterotrophs was statistically significant for one ecosystem (northern wooded savannah).

By contrast, at the maximum rate of harvesting (Figure 7b), significant changes in target cohort densities were seen in all ecosystems. Targeted omnivores declined by 84% in the desert ecosystem, 63%-75% in forest ecosystems, 50-64% in the wooded savannah, and around 20% in the grassland ecosystem. Densities of medium-large herbivores (3.2-32.6kg) declined, on average, by 53%-55% in the desert, 48%-52% in forest ecosystems, 43-53% in wooded savannah, and 40% in the grassland and shrub ecosystem. Despite being targeted for harvesting, small-bodied (1-3.2kg) herbivores were largely unaffected or even increased in abundance (in the grassland and shrub ecosystem). Targeted carnivores were largely unaffected (with the exception of the woodland and shrub ecosystem) though sample sizes were relatively small and outcomes had significant variation. Densities of non-target small-bodied (0.3-1kg) herbivores increased significantly in all ecosystems: by 161% in the wooded savannah in the South; 262% in the grasslands; 448%-648% in the forest ecosystems; and by over 1000% in the desert and the wooded savannah in the North. Small-bodied omnivore densities were also expected to increase: by between 39%-84% in the wooded savannah and forest ecosystems and by 268% on average in the grassland ecosystem.

Annual harvest of 90% of small and medium-sized heterotrophs (Figure 8) resulted in catastrophic declines in the target group densities in all ecosystems, losing 96% of herbivores and 99% of omnivores in the desert ecosystem; 88% of herbivores and 94% of omnivores in the tropical forest; 65% of herbivores and 95% of omnivores in the woodland and shrub; 74% of herbivores and 87% of omnivores in the wooded savannah ecosystems; and 41% of herbivores and 81% of omnivores in the grassland and shrub. Within the target group, smaller-bodied (1-3.2kg) herbivores were more resilient to harvesting than medium and large-bodied herbivores and omnivores. Densities of small-bodied non-target herbivores (0.3-1kg) increased by approximately 300% in the wooded savannah in the South, by over 4000% in the tropical forest, and by almost 9000% in the northern wooded savannah.

## 4 Discussion

The purpose of employing the Madingley Model here was to explore how potential bushmeat yields, maximum harvesting rates, and the impact of harvesting, might vary across African ecosystems. The model predicted that potential bushmeat yields varied by a factor of ten (or factor of three if we ignore desert). The harvesting rates required to achieve these yields did not vary significantly (55% to 65% per year, except for desert at 35%). The impact on ecosystem structure of harvesting at the maximum rates (harvest rates that maximised yield) varied quantitatively, but the qualitative pattern was relatively consistent (Figure 7 and Figure 8). Results such as these, produced by general ecosystem models, which are in their infancy, should be treated with caution (Purves *et al.*, 2013; Harfoot *et al.*, 2014). However, this class of models is at least able to begin to explore questions for which direct data are currently almost entirely lacking (Travers *et al.*, 2007; Link, Fulton and Gamble, 2010; Bartlett *et al.*, 2016; Newbold *et al.*, 2018).

A thorough mathematical analysis, outside the remit of the paper, would be needed to understand exactly why the Madingley Model made the predictions it did for potential yields from bushmeat hunting in Africa. No other variability has been introduced to the model’s inputs, except for the variation in the ecosystems’ structure and function which emerge in the model by varying the location of harvesting simulations, and it is possible that important variation in ecosystem parameters has been missed. Nonetheless there is sufficient evidence to make two tentative conclusions.

First, it is notable that animal biomasses (and therefore the potential bushmeat yields) are not predicted simply by Net Primary Productivity (NPP) (Lieth, 1975; Coe *et al*., 1976; Levin, 1998). NPP, which measures the total annual production of plant material (Roxburgh *et al.*, 2005) and is the ultimate source of productivity for all other ecosystem components including the animals targeted in bushmeat hunting (Del Grosso *et al.*, 2008; Petz *et al.*, 2014), is greatest in the tropical forest (Kicklighter *et al.*, 1999), whereas the greatest potential bushmeat yields appear in savannahs and woodlands. This simple result suggests that the potential bushmeat yields reflect the overall structure and function of the ecosystem, which emerges from a complex interaction between climate, plants, and animals, in a way which is at least partly, and approximately, captured by the Madingley Model.

Second, the potential yields were greatest where the ecosystem in the pristine state had higher total biomass represented in functional groups targeted by the bushmeat hunting. Higher biomasses of endothermic small and medium-sized (1-23kg) heterotrophs were returned in the wooded savannah ecosystems than in the forests, grasslands and shrub, and the southern desert, with the latter two ecosystems dominated by large-bodied herbivores outside our harvesting target range (Figure 4). Although empirical estimates of bushmeat yields were not available for the majority of the modelled ecosystems (though see below regarding yield estimates in the tropical forests of the Congo Basin), the ecosystems biomass pyramids (Figure 4) corresponded relatively well with the current literature (Bell, 1982; Bennett and Robinson, 2000). For example, high biomasses of large-bodied herbivores in arid and semi-arid ecosystems (southern desert, grasslands and wooded savannah), and low herbivore biomasses in the forest ecosystems, corresponded with Bennett and Robinson’s (2000) estimates of high mammalian biomasses in the open grasslands and woodlands (5-7 times higher than the evergreen forest) and low abundances of ungulates in tropical forests (attributed to the scarcity of grasses and browse) (Table 3). Similarly, high total biomasses of small and medium-sized herbivores in the grasslands and the northern wooded savannah agreed with Bell’s (1982) estimates of high densities of small herbivores in open short- and medium-length grasslands of East-African savannahs.

**Table 3.** Comparison of the Madingley Model’s estimates of animal biomasses (adult body mass≥1kg; with no harvesting), vs observed animal biomasses of mammals (body weight≥1kg) in sub-Saharan Africa (Bennett and Robinson, 2000).

| Ecosystem | Total animal biomass (kg/km<sup>2</sup>) |  |
| --- | --- | --- |
|  | Model Outputs | Observed for mammals |
| Evergreen forests | 150000 | >3000 |
| Open forests/grasslands | 170000-200000 | circa 15000 |
| Open grasslands | 240000 | circa 20000 |

However, the number of inverted trophic pyramids in our results (Figure 3) was surprising (Elton, 1927). Trebilco *et al*. (2013) showed that top-heavy pyramids could indicate an overestimation of predator abundance or energy available to carnivores. On the other hand, the Madingley Model predictions are for pristine terrestrial ecosytems, for which very little data on trophic pyramids are available. The model also predicted the carnivores to be predomonantly ectothermic (Figure 4), which is consistent with the known evolutionary history of ectotherms (ectothermic top carnivores were believed to be 5 times heavier than endothermic top carnivores; Burness *et al*., 2002), but is not consistent with these ecosystems today. This disparity is likely to indicate a problem in the formulation of one or mode ecological processes in the Madingley Model, the diagnosing and correction of which is outside the scope of this paper. The possible overestimation of the abundance of large-bodied carnivores, and the overestimation of the proportion of these carnivores that are ectothermic, together explain the very high biomass estimates for ectothermic carnivores predicted here. High biomass estimates for large-bodied carnivores in the more productive forest and savannah ecosystems in the Model were also reported by Harfoot *et al.* (2014).

The Madingley Model predicted bushmeat yields that were substantial on a per capita basis (Table 1). However, the model also predicted that bushmeat harvesting at these rates would have profound effects on ecosystem structure, with substantial reductions in target functional groups (reductions of 80% or more were typical; Table 2) coupled with substantial increases in non-target groups (increases of 200% or more were typical; Figure 7). These effects were not restricted to just one, sensitive ecosystem, but seen across all of the ecosystems. Such large ecosystem impacts call for a careful consideration of what it means for a harvest policy to be deemed sustainable (see below). Further work could examine the potential impacts of shifting the harvesting in response to the local biomass pyramid. For example, it would make sense to harvest larger animals in the savannahs, compared to size classes harvested here, which were based on bushmeat hunting data mainly from forest ecosystems (Fa, Ryan and Bell, 2005).

The quantitative ecosystem impacts of harvesting differed among the ecosystems, something that may not be obvious at first when viewing the summary figures (Figure 7 and Figure 8). For example, overall, the northern and southern savannahs showed similar impacts from harvesting (Figure 7); however, the northern savannah showed a large (circa 20 times) increase in small-bodied herbivorous endotherms not seen in the southern savannah (circa 2 times increase only). The grassland and woodland ecosystems had the highest extinction rates without harvesting (Figure 5). The exact reasons for high background extinction rates in the woodland and shrub and the grasslands ecosystems (Figure 5) are unclear and could be addressed in future work. One possible explanation could be a higher share of smaller-bodied animals with shorter life spans and higher rates of turnover compared to other ecosystems (although based on Figure 4, this was not the case). Opposite to expectation (Woodroffe, 2000; Azhar *et al.*, 2014; Newbold *et al.*, 2018), the omnivores were more sensitive to harvesting than the carnivores and herbivores. The omnivores had the lowest total biomass in all simulated ecosystems except for the deserts (Figure 3) with a higher share of medium-sized animals compared to the other functional groups (Figure 4). The non-linear responses to exploitation are a manifestation of complex trophic interactions and dynamic predator-prey responses in the Madingley Model (Newbold *et al.*, 2018). The omnivores’ higher sensitivity to harvesting could be explained by a combination of harvesting, increased competition for limited resources and an increase in predation (Figure 7 and Figure 8).

The 90% removal of all animals simulated here (Figure 8) is not likely in real-life systems; nevertheless, the model results show that such intensive harvesting would have profound effects on ecosystem structure. Empirical evidence of ecosystem responses to perturbations is still limited (Newbold *et al.*, 2018) with studies focusing on particular ecosystems and on incomplete subsets of the species in these ecosystems (though see Frank *et al.*, 2005; Carpenter *et al.*, 2011). These results underscore the need for ecosystem-specific studies to inform harvesting policies. Overall, grasslands and wooded savannah were the least affected by harvesting, and tropical forest and deserts the most affected. A global analysis of variances in vegetation productivity over the past 14 years identified tropical forests and desert regions of Africa as more sensitive to climate variability compared to savannah regions, which suggested that these areas were also more sensitive to anthropogenic pressures (Seddon *et al.*, 2016), such as bushmeat harvesting.

The low impact of harvesting on carnivore abundances was explained by a very low percentage of endothermic target carnivores in the pristine state in all ecosystems (below 1% of total biomass, with the exception of the tropical forest). The variation in predicted impacts of harvesting on carnivore abundances was high (Figure 7 and Figure 8), and any potential impacts of harvesting on carnivore abundances may have been offset (fully or partially) by large increases in abundance of their small-bodied prey. Nevertheless, for this region, the Madingley Model appears to have a structural problem with this aspect of its predictions – although good data is lacking, it is impossible to believe that over 90% of mid- and large-sized carnivores in these ecosystems are ectothermic (or would be, in the pristine state that is being simulated). This problem does not necessarily have a large overall impact on the Madingley Model used for general questions, but it is of central importance here because the harvesting policy distinguishes between these two groups. Complete absence of carnivores in some of the simulated ecosystems (e.g. in desert ecosystem, also reported by Newbold *et al.*, 2018) is also unrealistic. Further work could seek to improve the model, and in the meantime, examine the predicted yields if the ectotherms were effectively treated as endotherms for the purposes of hunting removals.

The model’s predictions for potential bushmeat yields were large enough to have implications for human nutrition. When taking human population density into account, the annual yield per capita was 67 kg for northern grass and shrub; over 200 kg for northern and southern wooded savannah, and tropical forest (desert was an exception, given the lack of human population data). To put these figures into context, the annual meat consumption per capita in the United States is estimated to be 62 kg (FAO, 2013), although a fairer comparison is with US meat production, at 124 kg (losses between production and consumption are around 50%).

Are these predictions realistic? Data are scarce, but the model’s estimate of yields under the high risk strategy for the tropical forest ecosystem of 2246.98 kg km^−2^ year^−1^ (±36.89) compared surprisingly well with estimated meat offtake in the Congo basin. The model’s estimated yields were within an order of magnitude of 645 kg km^−2^ year^−1^ bushmeat offtakes reported by Wilkie and Carpenter (1999) for the Congo Basin, and overlapped with the Congo Basin estimates by Fa, Ryan and Bell (2005) of 2645 kg km^−2^ year^−1^. Taken at face value, then, the Madingley Model predicts that this rate of hunting is sustainable, at least within this ecosystem, and suggests further that even higher sustainable yields are possible in savannahs. However, there are several important caveats here. First, as mentioned above, according to the model, harvesting at these rates has drastic effects on ecosystem structure, and so is sustainable in the narrow sense only. Second, again according to the model, to achieve the maximum yield in the tropical rainforest requires the removal of 30% of all animals in the target group (i.e. all carnivores, herbivores and omnivores of body mass 1-23 kg) every year. This may not be feasible in practice (e.g. due to logistical constraints), and even if it was, underscores why such harvesting would be likely to have profound effects on the ecosystem. The final caveat is a reminder that general ecosystem models, such as the Madingley, are still in their infancy, and as such their predictions should be treated with caution. Nonetheless, the results do suggest that substantial sustainable bushmeat yields may be possible in African ecosystems – and that general ecosystem models can begin to estimate these yields, and/or raise important questions for further study.

The differences between predicted bushmeat yields and reported beef offtakes (Otte and Chilonda, 2002) were higher in the tropical forest and wooded savannah ecosystems, and relatively low in the grasslands, where predicted bushmeat yields were at the their lowest and beef offtakes were maximised (Table 1). Cattle, goats, sheep, poultry and pigs all contribute to protein intake in Africa; however, the livestock distribution across Africa is uneven, with more than half of all ruminant livestock in sub-Saharan Africa concentrated in the arid and semi-arid areas (Otte and Chilonda, 2002). Intensive land management including animal husbandry has been shown to significantly impact biodiversity, particularly in pristine ecosystems (Newbold *et al.*, 2015). If achieved, sustainable well-regulated bushmeat harvesting could help alleviate some of the negative impacts of livestock husbandry by providing an alternative source of protein in the tropical forests of Africa, at least in the near future.

Because the model was set to target small and medium-sized animals, it did not necessarily capture the highest possible yields in each ecosystem. The decision to keep the body size of the target group constant was based on: a) the sizes of animals caught by snare, bow and arrow, or rifle, by a single hunter (Fa, Peres and Meeuwig, 2002); b) the complexities of identifying animal sizes that maximised yields in each ecosystem: these ‘optimal’ animal sizes may or may not be reasonable in reality, and c) the ease of comparison between ecosystems. One could also argue that the preference for small and medium-sized animals was more conservative, due to lower reproductive rates and densities of larger-bodied animals. The question of optimal body sizes for harvesting in different ecosystems can be explored in future work.

By harvesting once a year (rather than continuously) and assuming constant, non-adaptive harvesting we might have disadvantaged ecosystems with higher seasonality (such as grasslands). More sophisticated harvesting strategies could be implemented, though one could argue that more sophisticated harvesting regimes would make the modelled processes more obscure and could confound interpretation of the results.

Here, we examined how the ecosystems differed in their capacity to support bushmeat harvesting and in responses to harvesting, as predicted by the Madingley Model. Although it wasn’t possible to identify the exact ecological interactions and processes that determined ecosystems capacity for supporting sustainable bushmeat yields, some ecosystems were much more productive and resilient to harvesting than the others suggesting that the ecosystem structure and functioning were important predictors of productivity and resilience. Because the Madingley Model does not require specific parameter inputs (Harfoot *et al.*, 2014), we were able to compare the dynamics of ecosystem communities consisting of species that we may not have population parameter estimates for, and therefore, may not be able to model otherwise. In addition, the modelled ecosystem communities not only incorporated the effects of multi-trophic interactions but also the effects of the environmental conditions on plant and animal biomasses. As climate plays a crucial role in determining ecosystem features (e.g. Coe *et al*., 1976; Levin, 1998), it follows that the ecosystems capacity for wild meat production, as well as livestock husbandry, will change in the future. These results are experimental, but they demonstrate the potential of a general ecosystem model such as the Madingley Model, as an additional tool for informing decisions in conservation and land management.

## Acknowledgements

This work was part of a PhD thesis funded by the Natural Environment Research Council (TB). We thank Mike Harfoot for valuable discussions about this work, and Tim Newbold and Luca Borger for comments on the manuscript.

## References

Azhar, B., Lindenmayer, D.B., Wood, J., Fischer, J. and Zakaria, M. (2014) ‘Ecological impacts of oil palm agriculture on forest mammals in plantation estates and smallholdings’, Biodiversity and Conservation, 23(5), pp. 1175–1191. doi: 10.1007/s10531-014-0656-z.

Barnes, R. F. W. and Lahm, S. A. (1997) ‘An Ecological Perspective on Human Densities in the Central African Forest’, Journal of Applied Ecology. [British Ecological Society, Wiley], 34(1), pp. 245–260. doi: 10.2307/2404862.

Bartlett, L. J. Newbold, T., Purves, D.W., Tittensor, D.P. and Harfoot, M.B.J. (2016) ‘Synergistic impacts of habitat loss and fragmentation on model ecosystems’, in Proc. R. Soc. B. The Royal Society, p. 20161027.

Bell, R. H. V (1982) ‘The effect of soil nutrient availability on community structure in African ecosystems’, in Ecology of tropical savannas. Springer, pp. 193–216.

Bennett, E. L. and Robinson, J. G. (2000) ‘Carrying capacity limits to sustainable hunting in tropical forests’, Hunting for sustainability in tropical forests, pp. 13–30.

Boote, K. J., Jones, J. W. and Pickering, N. B. (1996) ‘Potential uses and limitations of crop models’, Agronomy journal. American Society of Agronomy, 88(5), pp. 704–716.

Borrett, S. R., Moody, J. and Edelmann, A. (2014) ‘The rise of Network Ecology: Maps of the topic diversity and scientific collaboration’, Ecological Modelling, 293, pp. 111–127. doi: https://doi.org/10.1016/j.ecolmodel.2014.02.019.

Burness, G., Diamond, J. and Flannery, T. (2002) Dinosaurs, dragons, and dwarfs: The evolution of maximal body size, Proceedings of the National Academy of Sciences of the United States of America. doi: 10.1073/pnas.251548698.

Butt, N., Malhi, Y., Phillips, O. and New, M. (2008) ‘Floristic and Functional Affiliations of Woody Plants with Climate in Western Amazonia’, Journal of Biogeography. Wiley, 35(5), pp. 939–950. Available at: http://www.jstor.org/stable/30142883.

Carpenter, S. R., Cole, J.J., Pace, M.L., Batt, R., Brock, W.A., Cline, T., Coloso, J., Hodgson, J.R., Kitchell, J.F., Seekell, D.A., Smith, L. and Weidel, B. (2011) ‘Early warnings of regime shifts: a whole-ecosystem experiment’, Science. American Association for the Advancement of Science, 332(6033), pp. 1079–1082.

Christensen, V. and Walters, C. J. (2004) ‘Ecopath with Ecosim: methods, capabilities and limitations’, Ecological Modelling, 172(2), pp. 109–139. doi: https://doi.org/10.1016/j.ecolmodel.2003.09.003.

Coe, M. J., Cumming, D. H. and Phillipson, J. (1976) ‘Biomass and production of large African herbivores in relation to rainfall and primary production’, Oecologia. Springer, 22(4), pp. 341–354.

Crête (2001) ‘The distribution of deer biomass in North America supports the hypothesis of exploitation ecosystems’, Ecology Letters. John Wiley & Sons, Ltd, 2(4), pp. 223–227. doi: 10.1046/j.1461-0248.1999.00076.x.

Elton, C. S. (Charles S. (1927) Animal ecology /C.S. Elton. London: London: Sidgwick and Jackson.

Fa, J. E., Ryan, S. F. and Bell, D. J. (2005) ‘Hunting vulnerability, ecological characteristics and harvest rates of bushmeat species in afrotropical forests’, Biological conservation. Elsevier, 121(2), pp. 167–176.

Flores, C., Kortsch, S., Tittensor, D., Harfoot, M., Purves, D. (2019) ‘Food Webs: Insights from a General Ecosystem Model’, BioRxiv. Cold Spring Harbor Laboratory, p. 588665.

Frank, K. T., Petrie, B., Choi, J.S. and Leggett, W.C. (2005) ‘Trophic cascades in a formerly cod-dominated ecosystem’, Science. American Association for the Advancement of Science, 308(5728), pp. 1621–1623.

Fulton, E.A., Link, J.S., Kaplan, I.C., Savina-Rolland, M., Johnson, P., Ainsworth, C., Horne, P., Gorton, R., Gamble, R.J., Smith, A.D. and Smith, D.C. (2011) ‘Lessons in modelling and management of marine ecosystems: the Atlantis experience’, Fish and Fisheries. Wiley Online Library, 12(2), pp. 171–188.

Del Grosso, S., Parton, W., Stohlgren, T., Zheng, D., Bachelet, D., Prince, S., Hibbard, K. and Olson, R. (2008) ‘Global potential Net Primary Production predicted from vegetation class, precipitation, and temperature’, Ecology. Wiley-Blackwell, 89(8), pp. 2117–2126. doi: 10.1890/07-0850.1.

Harfoot, M.B., Newbold, T., Tittensor, D.P., Emmott, S., Hutton, J., Lyutsarev, V., Smith, M.J., Scharlemann, J.P. and Purves, D.W. (2014) ‘Emergent global patterns of ecosystem structure and function from a mechanistic general ecosystem model.’, PLoS biology. Edited by M. Loreau. Public Library of Science, 12(4), p. e1001841. doi: 10.1371/journal.pbio.1001841.

Hirota, M., Holmgren, M., Van Nes, E. H. and Scheffer, M. (2011) ‘Global Resilience of Tropical Forest and Savanna to Critical Transitions’, Science, 334(6053), pp. 232–235. Available at: http://science.sciencemag.org/content/334/6053/232.abstract.

Hunter, M. D. and Price, P. W. (1992) ‘Playing Chutes and Ladders: Heterogeneity and the Relative Roles of Bottom-Up and Top-Down Forces in Natural Communities’, Ecology. Ecological Society of America, 73(3), pp. 724–732. Available at: http://www.jstor.org/stable/1940152.

Kicklighter, D. W., Bondeau, A., Schloss, A.L., Kaduk, J. and Mcguire, A.D. (1999) ‘Comparing global models of terrestrial net primary productivity (NPP): global pattern and differentiation by major biomes’, Global Change Biology. John Wiley & Sons, Ltd (10.1111), 5(S1), pp. 16–24. doi: 10.1046/j.1365-2486.1999.00003.x.

Korzukhin, M. D., Ter-Mikaelian, M. T. and Wagner, R. G. (1996) ‘Process versus empirical models: which approach for forest ecosystem management?’, Canadian Journal of Forest Research. NRC Research Press, 26(5), pp. 879–887.

Krinner, G., Viovy, N., de Noblet-Ducoudré, N., Ogée, J., Polcher, J., Friedlingstein, P., Ciais, P., Sitch, S. and Prentice, I.C. (2005) ‘A dynamic global vegetation model for studies of the coupled atmosphere-biosphere system’, Global Biogeochemical Cycles. Wiley Online Library, 19(1).

Levin, S. A. (1998) ‘Ecosystems and the biosphere as complex adaptive systems’, Ecosystems. Springer, 1(5), pp. 431–436.

Lieth, H. (1975) ‘Modeling the Primary Productivity of the World BT - Primary Productivity of the Biosphere’, in Lieth, H. and Whittaker, R. H. (eds). Berlin, Heidelberg: Springer Berlin Heidelberg, pp. 237–263. doi: 10.1007/978-3-642-80913-2_12.

Link, J. S., Fulton, E. A. and Gamble, R. J. (2010) ‘The northeast US application of ATLANTIS: A full system model exploring marine ecosystem dynamics in a living marine resource management context’, Progress in Oceanography. Pergamon, 87(1–4), pp. 214–234. doi: 10.1016/J.POCEAN.2010.09.020.

Mace, G. M. and Lande, R. (1991) ‘Assessing extinction threats: toward a reevaluation of IUCN threatened species categories’, Conservation Biology. Wiley Online Library, 5(2), pp. 148–157.

McNaughton, S. J. (1976) ‘Serengeti Migratory Wildebeest: Facilitation of Energy Flow by Grazing’, Science. American Association for the Advancement of Science, 191(4222), pp. 92–94. Available at: http://www.jstor.org/stable/1741873.

Mokany, K., Ferrier, S., Connolly, S.R., Dunstan, P.K., Fulton, E.A., Harfoot, M.B., Harwood, T.D., Richardson, A.J., Roxburgh, S.H., Scharlemann, J.P.W., Tittensor, D.P., Westcott, D.A. and Wintle, B.A. (2016) ‘Integrating modelling of biodiversity composition and ecosystem function’, Oikos. Wiley Online Library, 125(1), pp. 10–19.

Montoya, J. M., Pimm, S. L. and Solé, R. V (2006) ‘Ecological networks and their fragility’, Nature, 442(7100), p. 259. doi: 10.1038/nature04927.

Newbold, T., Hudson, L.N., Hill, S.L., Contu, S., Lysenko, I., Senior, R.A., Börger, L., Bennett, D.J., Choimes, A., Collen, B. and Day, J. (2015) ‘Global effects of land use on local terrestrial biodiversity’, Nature. Nature Publishing Group, a division of Macmillan Publishers Limited. All Rights Reserved., 520, p. 45. Available at: https://doi.org/10.1038/nature14324.

Newbold, T., Tittensor, D.P., Harfoot, M.B., Scharlemann, J.P.W and Purves, D.W. (2018) ‘Non-linear changes in modelled terrestrial ecosystems subjected to perturbations’, bioRxiv, p. 439059. doi: 10.1101/439059.

Otte, M. J. and Chilonda, P. (2002) ‘Cattle and small ruminant production systems in sub-Saharan Africa. A systematic review’. Rome (Italy) FAO.

Parrott, L. and Meyer, W. S. (2012) ‘Future landscapes: managing within complexity’, Frontiers in Ecology and the Environment. Wiley Online Library, 10(7), pp. 382–389.

Peres, C. A. (2000) ‘Effects of subsistence hunting on vertebrate community structure in Amazonian forests’, Conservation Biology. Wiley Online Library, 14(1), pp. 240–253.

Petz, K., Alkemade, R., Bakkenes, M., Schulp, C.J.E., van der Velde, M. and Leemans, R. (2014) ‘Mapping and modelling trade-offs and synergies between grazing intensity and ecosystem services in rangelands using global-scale datasets and models’, Global Environmental Change, 29, pp. 223–234. doi: https://doi.org/10.1016/j.gloenvcha.2014.08.007.

Purves, D., Scharlemann, J.P.W., Harfoot, M., Newbold, T, Tittensor, D.P., Hutton, J. and Emmott, S. (2013) ‘Time to model all life on Earth’, Nature. Nature Publishing Group, a division of Macmillan Publishers Limited. All Rights Reserved., 493, p. 295. Available at: http://dx.doi.org/10.1038/493295a.

Roxburgh, S. H., Berry, S.L., Buckley, T.N., Barnes, B. and Roderick, M.L. (2005) ‘What is NPP? Inconsistent accounting of respiratory fluxes in the definition of net primary production’, Functional Ecology. Wiley Online Library, 19(3), pp. 378–382.

Seddon, A., Macias-Fauria, M., Long, P., Benz, D. and Willis, K. (2016) ‘Sensitivity of global terrestrial ecosystems to climate variability’, Nature. London, 531(7593), pp. 229–232K. doi: 10.1038/nature16986.

Sol, R. V and Montoya, M. (2001) ‘Complexity and fragility in ecological networks’, Proceedings of the Royal Society B: Biological Sciences, 268(1480), pp. 2039–2045. doi: 10.1098/rspb.2001.1767.

Stevens, B. M., Propster, J., Wilson, G.W.T., Abraham, A., Ridenour, C., Doughty, C. and Johnson, N.C. (2018) ‘Mycorrhizal symbioses influence the trophic structure of the Serengeti’, journal of ecology. Wiley Online Library, 106(2), pp. 536–546.

Thompson, R. M., Brose, U., Dunne, J. A., Hall Jr, R. O., Hladyz, S., Kitching, R. L., Martinez, N.D., Rantala, H., Romanuk, T.N., Stouffer, D.B. and Tylianakis, J.M. (2012) ‘Food webs: reconciling the structure and function of biodiversity’, Trends in ecology & evolution. Elsevier, 27(12), pp. 689–697.

Travers, M., Shin, Y.-J., Jennings, S. and Cury, P. (2007) ‘Towards end-to-end models for investigating the effects of climate and fishing in marine ecosystems’, Progress in oceanography. Elsevier, 75(4), pp. 751–770.

Woodroffe, R. (2000) ‘Predators and people: using human densities to interpret declines of large carnivores’, Animal Conservation. 2000/05/01. Cambridge University Press, 3(2), pp. 165–173. doi: DOI: undefined.

Yates, D. N., Kittel, T. G. F. and Cannon, R. F. (2000) ‘Comparing the correlative Holdridge model to mechanistic biogeographical models for assessing vegetation distribution response to climatic change’, Climatic Change. Springer, 44(1–2), pp. 59–87.

