## Supplemental Table 1 for "Modelling variation in bushmeat harvesting among seven African ecosystems using the Madingley Model: yield, survival and ecosystem impacts"

**Appendix 1 Geographic coordinates of the Madingley harvesting simulations.**

| Location | Vegetation Type | Coordinates |
| --- | --- | --- |
| Desert and desert shrub – North | Desert | 19^0^N 22^0^W |
| Grass and Shrub – North | Savannah | 10^0^N 22^0^W |
| Wooded Savanna – North | Savannah | 7^0^N 22^0^W |
| Tropical Forest | Forest | 0^0^N 22^0^W |
| Woodland and Shrub – South | Forest | 9^0^S 22^0^W |
| Wooded Savanna – South | Savannah | 16^0^S 22^0^W |
| Desert and Desert Shrub – South | Desert | 30^0^S 22^0^W |
