## Supplemental Figure 2 for "Modelling variation in bushmeat harvesting among seven African ecosystems using the Madingley Model: yield, survival and ecosystem impacts"

**Appendix 2 Median densities of target species (with 95% confidence intervals) with annual harvest rate, by ecosystem.**

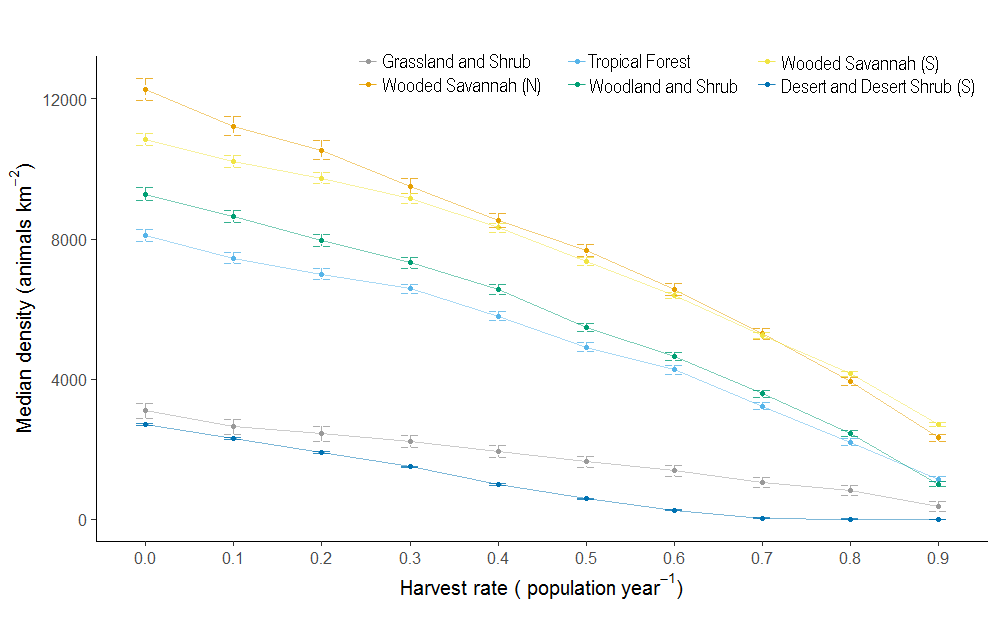
