## Supplemental Table 3 for "Modelling variation in bushmeat harvesting among seven African ecosystems using the Madingley Model: yield, survival and ecosystem impacts"

**Appendix 3 Average declines in target animal densities per 0.05 population year^-1^ increase in harvest rate,** $\boldsymbol{\varphi}$ **up to** $\boldsymbol{\varphi\leq}$**0.70, and per 0.10 population year^-1^ increase in harvest rate,** $\boldsymbol{\varphi}$**, thereafter, by ecosystem.**

| **Ecosystem** | **Harvest rate** $\boldsymbol{\varphi}$ | |
| --- | --- | --- |
|  | $\boldsymbol{\leq}$**0.70** | **0.70-0.90** |
| Grass and Shrub – North | 0.04 | 0.12 |
| Wooded Savanna – North | 0.06 | 0.28 |
| Tropical Forest | 0.06 | 0.31 |
| Woodland and Shrub – South | 0.06 | 0.32 |
| Wooded Savanna – South | 0.05 | 0.26 |
| Desert and Desert Shrub – South | 0.16 | 0.37 |
